# CSF infusion of TrkB agonist, 7,8-dihydroxyflavone, is ineffective in promoting remyelination in cuprizone and EAE models of multiple sclerosis

**DOI:** 10.1101/2021.04.19.440405

**Authors:** Jessica L Fletcher, Rhiannon J Wood, Alexa R Prawdiuk, Ryan O’Rafferty, Ophelia Ehrlich, David G Gonsalvez, Simon S Murray

## Abstract

Small molecular weight functional mimetics of brain-derived neurotrophic factor (BDNF) which act via the TrkB receptor have been developed to overcome the pharmacokinetic limitations of BDNF as a therapeutic for neurological disease. Activation of TrkB on oligodendrocytes has been identified as a potential strategy for myelin repair in demyelinating conditions. Here, we tested the efficacy of intracerebroventricular infusion of TrkB agonist 7,8-dihydroxyflavone (DHF) to promote myelin repair in the cuprizone model and alter the course of experimental autoimmune encephalomyelitis (EAE). In these two distinct mouse models used for the preclinical testing of remyelinating therapeutics, we found that DHF infusion increased the percentage of myelin basic protein and density of oligodendrocyte progenitor cells (OPCs) in the corpus callosum of female C57BL/6 mice after cuprizone demyelination. However, DHF did not alter the percentage of axons myelinated or increase the density of post-mitotic oligodendrocytes in this model. Direct cerebrospinal fluid infusion of DHF also had no effect on the clinical course of EAE, and examination of the lumbar spinal cord after 21 days of treatment revealed extensive demyelination. These results indicate that direct cerebrospinal fluid infusion of DHF is ineffective at promoting myelin repair in toxin-induced and inflammatory models of demyelination.

## Introduction

Neurotrophins have long been identified as therapeutic candidates to treat neurologic disorders due to their ability to promote neuronal survival and differentiation as well as synaptic plasticity [1]. However, it is only comparatively recently that the effects neurotrophins mediate *via* glial cells have received attention as a potential mechanism to prevent neurodegeneration [2,3]. This includes the effect of brain-derived neurotrophic factor (BDNF), which has been shown to promote central nervous system (CNS) myelination via its TrkB receptor [4,5]. That BDNF elicits oligodendroglial differentiation and myelination by activating TrkB is highly relevant for the auto-immune demyelinating disease, multiple sclerosis (MS). Although there are now many immune-targeted therapies that can significantly modify the course of relapsing-remitting MS, there are no treatments which can prevent or reverse the axon loss and neuron death that lead to the increasing severity of neurologic dysfunction experienced by patients with progressive MS [6,7]. Therefore, neuroprotective therapeutic strategies that can also promote myelin repair—such as selectively targeting the TrkB receptor—are highly sought-after as adjunct treatments to the immuno-modulatory therapies clinically available for MS.

Although attractive as a remyelinating therapeutic, BDNF itself has poor pharmacokinetic properties – it is a large molecule unable to cross the blood-brain barrier, that is rapidly cleared systemically [8] and is non-selective, interacting with both TrkB and the pan-neurotrophin receptor p75^NTR^ [1,2,9]. Collectively, these properties of BDNF have led to the development of multiple smaller molecular weight TrkB agonists, also called functional BDNF-mimetics [1,10–12]. This includes TDP6 and LM22A-4, both which promote myelin repair in an oligodendroglial TrkB-dependent manner in the cuprizone model of de- and remyelination, in which MS therapies for myelin repair are frequently tested [13,14]. Notably, 7,8-dihydroxyflavone (DHF), the most widely used TrkB agonist within the biomedical literature (Fig. S1) has not previously been tested in cuprizone. However, in experimental autoimmune encephalomyelitis (EAE) it has been shown to significantly reduce the course of clinical signs when administered prophylactically via intra-peritoneal injection [15]. In addition, DHF has undergone extensive biochemical and biophysical characterisation to demonstrate its ability to interact with the extracellular domain of TrkB and elicit TrkB phosphorylation *in vitro* [16]. As a flavonoid, DHF has also been shown to act as an anti-oxidant, promoting survival in neuronal cell culture models of oxidative stress independent of TrkB receptor expression [17,18]. Overall, the ability of DHF to elicit TrkB signalling and act as a neuroprotective anti-oxidant makes it a promising candidate to promote myelin repair in the context of MS and inflammatory demyelination.

To test the capacity of DHF to promote myelin repair in response to inflammatory demyelination, two mouse models of myelin loss were used, the cuprizone model and EAE. In the toxin-induced cuprizone model, mature oligodendrocytes are selectively killed resulting in profound demyelination and inflammatory gliosis, followed by rapid remyelination once cuprizone is withdrawn [13,14,19]. However, cuprizone does not mimic the auto-immune attack against myelin which characterises MS. This is achieved in EAE, in which mice are immunised against a myelin protein (typically myelin oligodendrocyte glycoprotein, MOG), which results in neuroinflammation and clinical signs of ascending paralysis, but determining if remyelination has occurred in EAE is challenging [20]. In the current study DHF was administered directly to the CNS via intracerebroventricular infusion into the cerebrospinal fluid in both models. This was done to enable comparison with previous cuprizone studies investigating the remyelinating potential of TrkB activation [13,14], and to limit modulation of the peripheral immune system in EAE. Additionally, in EAE, DHF was administered at the onset of clinical signs to reflect the likely experience of people with MS, who would receive treatment following initial clinical presentation.

## Results

### 7,8-dihydroxylfavone fails to enhance myelin repair in the cuprizone model of de- and remyelination

We have previously shown that 7-day ICV infusions of the structural peptide mimetic of BDNF, TDP6 and the partial TrkB agonist, LM22A-4 increase the level of MBP expression, the proportion of myelinated axons and myelin sheath thickness during remyelination after cuprizone-induced demyelination [13,14]. To determine if DHF could mediate the same effect as a TrkB agonist in the same context (Fig. 1A), we first assessed the level of MBP immunostaining in the caudal corpus callosum (Fig. 1C). This revealed 7-day infusion of DHF during remyelination resulted in an increased percentage area of MBP immunostaining compared to infusion with the aCSF vehicle (p=0.012), but not when compared to the 14% DMSO vehicle (p=0.22; Fig. 1B). To confirm whether this increase in MBP staining was due to a pro-myelinating effect, we next performed electron microscopy (EM; Fig. 1D), comparing the proportion of myelinated axons in the caudal corpus callosum of mice that received the aCSF vehicle, to those that received DHF, but did not detect a change (p=0.23, Fig. 1E). However, electron dense myelin debris was more frequently observed in the caudal callosum of mice that received DHF (Fig. 1D). Collectively, both the MBP and EM analyses indicate that 7-day ICV infusion of DHF itself does not enhance remyelination above either the infusion of aCSF or DMSO vehicle infusion.

**Figure 1:**
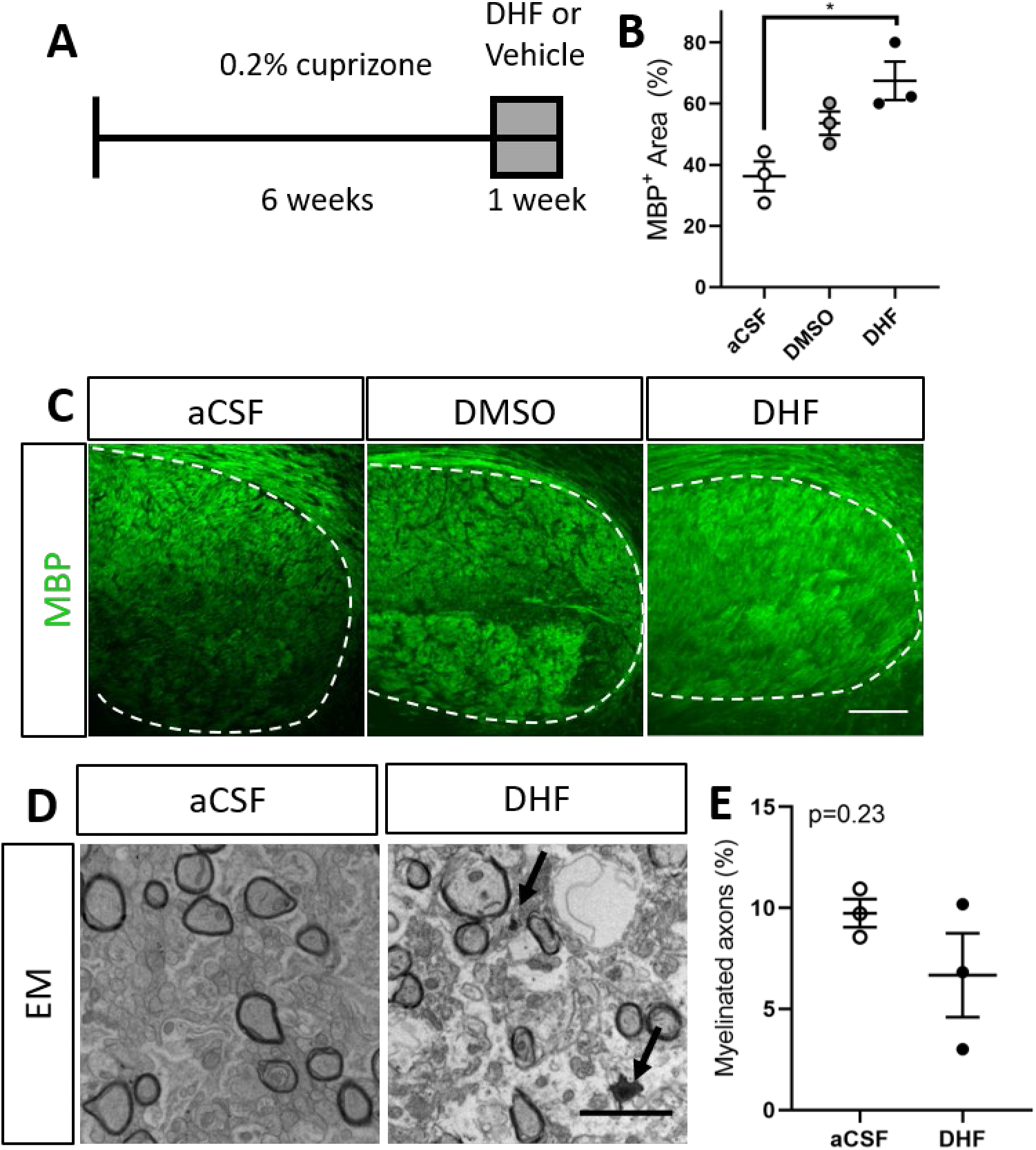
7,8-dihydroxylfavone increases area of myelin basic protein (MBP) immunostaining but does not increase the percentage of myelinated axons during remyelination. (a) Schematic of experimental paradigm, where cuprizone demyelinated mice received either DHF, aCSF or 14% DMSO vehicle *via* ICV infusion for 1 week. (b) Animals receiving DHF had an increased percentage area MBP^+^ immunostaining in the cadual corpus callosum, compared to mice that received the aCSF vehicle (p=0.012) but not compared to those that received the DMSO vehicle (p=0.22) (n=3/group, 1-way ANOVA, F_2,6_=0.09, Tukey’s post-hoc comparison). Representative (c) micrographs of MBP immunostaining (scale bar = 100μm) and (d) electron micrographs (scale bar = 2μm) in the caudal corpus callosum of aCSF, DMSO and DHF treated animals (arrows: myelin debris). (E) DHF treatment did not alter the percentage of axons myelinated in the caudal corpus callosum during remyelination, compared to animals that received aCSF (n=3/group, unpaired t-test, F_2,2_=9.04).

### 7,8-dihydroxyflavone increases OPC density after cuprizone-induced demyelination

Infusion of either TDP6 or LM22A-4 for 7 days during remyelination increased the density of total Olig2^+^ oligodendroglia in the caudal corpus callosum, primarily by increasing the density of Olig2^+^CC1^+^ post-mitotic oligodendrocytes [13,14]. Here, we sought to determine if DHF, mediated the same effect by performing cell counts on sagittal brain sections triple immunolabeled for Olig2 to identify all cells in the oligodendroglial lineage, PDGFRα to identify oligodendrocyte precursor cells (OPCs) and CC1 to identify post-mitotic oligodendrocytes (Fig. 2A). This identified a trend towards an increased density of Olig2^+^ oligodendroglia in the caudal corpus callosum of mice that received DHF treatment compared to those that received either aCSF or the DMSO vehicle (Fig. 2B). Treatment with DHF increased the density of OPCs (p=0.0002, Fig 2C), whereas DMSO infusion did not result in a detectable difference in OPC density in the caudal corpus callosum when compared to mice treated the aCSF vehicle (p=0.71, Fig. 2C). There was no detectable change (p=0.96) in the density of Olig2^+^CC1^+^ post-mitotic oligodendrocytes in the caudal corpus callosum of mice that received DHF, compared to those that received aCSF or DMSO (Fig 2D). These data indicate that ICV infusion of DHF, but not its DMSO vehicle during the remyelinating period selectively triggers OPC proliferation.

**Figure 2:**
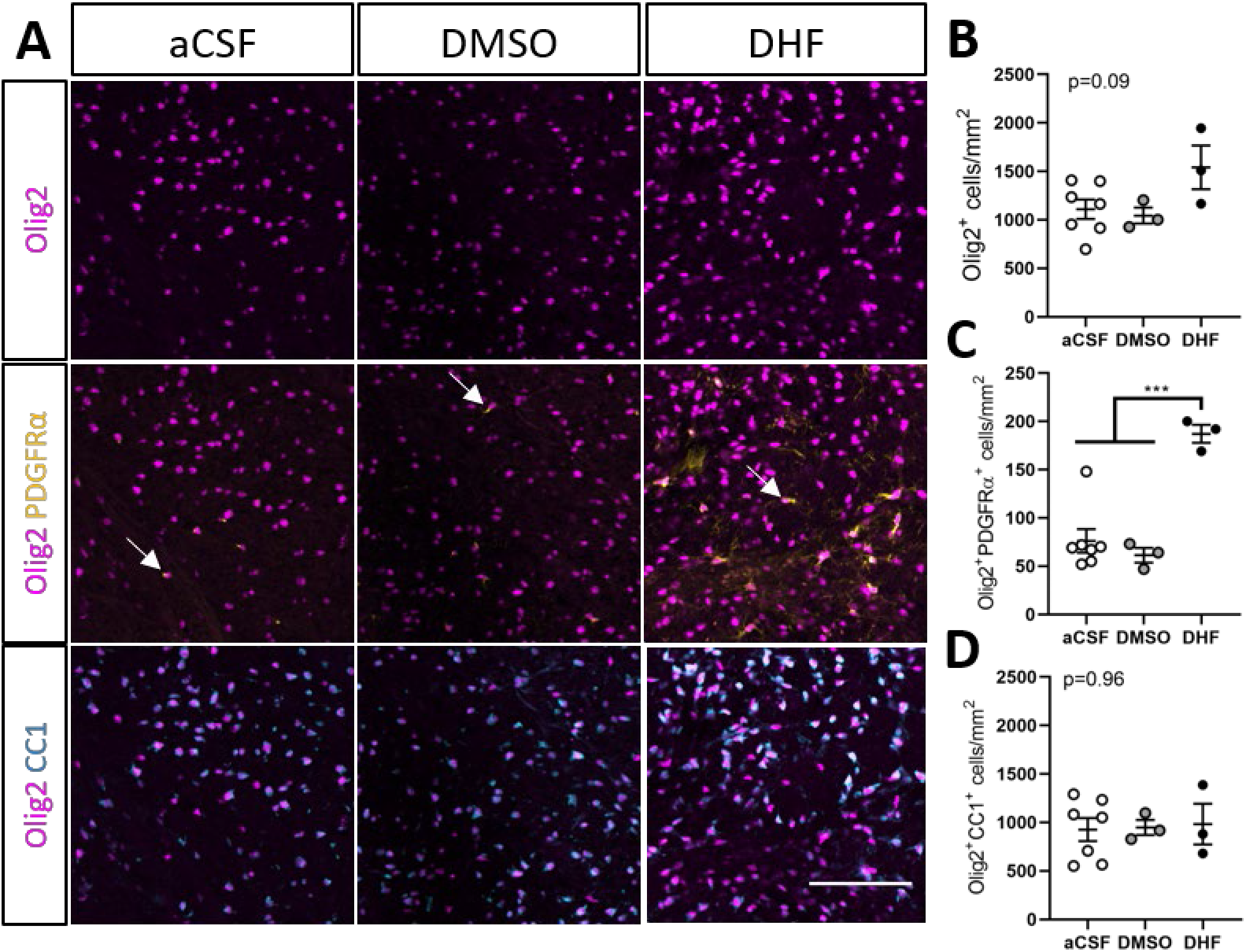
Increased oligodendrocyte progenitor cell density in the caudal corpus callosum of mice treated with 7,8-dihydroxylfavone during remyelination. (a) Representative micrographs for Olig2-PDGFRα-CC1 triple immunolabelling in the cadual corpus callosum of mice treated with aCSF, DMSO or DHF during remyelination. Scale bar=100μm. Brightness and contrast have been adjusted to improve visibility. (b) The density of Olig2+ oligodendroglia demonstrated a trend increase in mice that were treated with DHF compared to those that received the aCSF and DMSO vehicles. This was driven by (c) an increase (p=0.0002) in Olig2+PDGFRα+ OPC (arrows in A) densities in the corpus callosum of mice that received DHF compared to those that received aCSF or DMSO vehicles. (d) There was no change in the density of Olig2+CC1+ cells in the corpus callosum of mice treated with DHF compared to aCSF and DMSO. For (b-c) (*n*=3-7/group, 1-way ANOVA, Tukey’s post-hoc comparison).

### 7,8-dihydroxyflavone administered directly into the CNS after onset of clinical signs does not alter the clinical course of EAE

Others have shown that daily intraperitoneal injection of DHF in DMSO from the day of EAE induction can improve its clinical course and improve neuropathological outcomes including the extent of inflammation and demyelination [15]. We sought to build on these findings to determine if DHF treatment had a direct effect on the CNS during EAE via ICV infusion, and if it could alter the course of EAE after the onset of clinical signs (Fig. 3A). We found that compared to the treatment with DMSO vehicle, ICV infusion of DHF from Day 13 when mice exhibited tail weakness and/or weakness of one hindlimb, did not alter the clinical course of EAE (Treatment: p=0.47, Fig. 3B). We also observed that male mice induced with EAE had a tended to have more severe phenotype than female mice (Interaction Sex × Day: p=0.052, data not shown) but were underpowered to determine if DHF treatment altered this. Consistent with no effect of DHF on clinical score, we also found that compared to sham EAE animals (Fig. 3C), there remained profound demyelination in the lumbar spinal cord of DHF-treated EAE at Day 35, using SCoRe and Fluromyelin to detect compact myelin and myelin debris (Fig. 3C). Overall, these results indicate that direct infusion of DHF into the CNS after the onset of clinical signs, is unable to alter the course of EAE.

**Figure 3:**
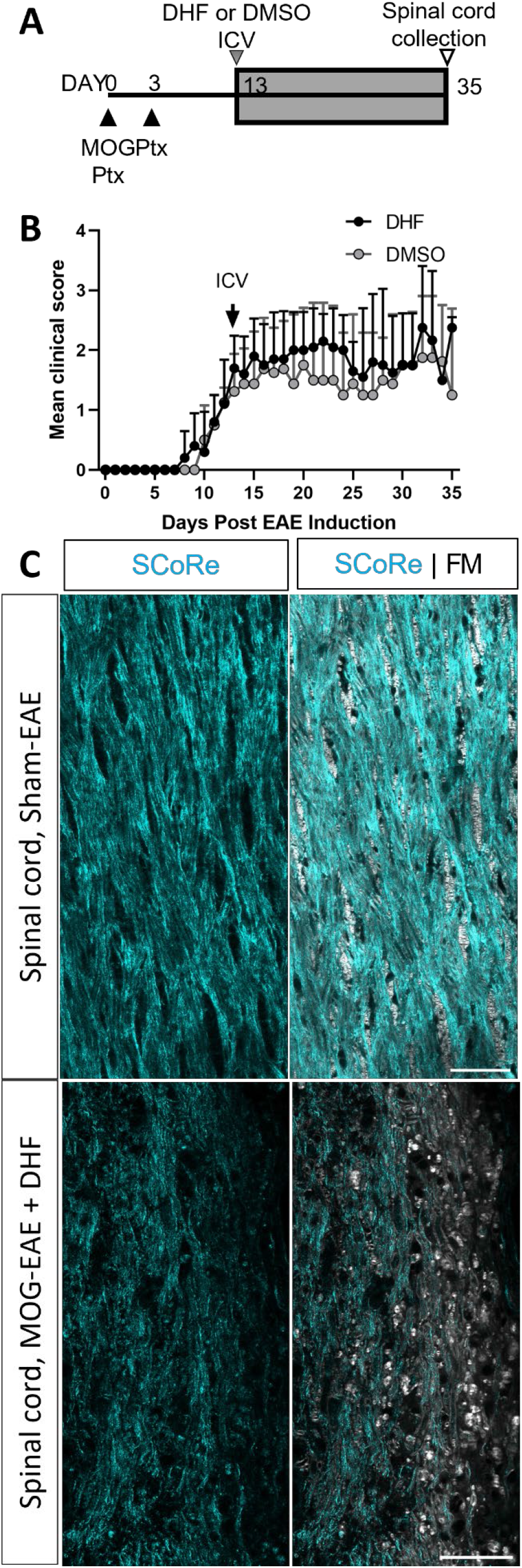
Direct CSF infusion of 7,8-dihydroxyflavone after the onset of clinical signs does not alter the disease course or restore myelin in EAE. (a) Schematic of experimental paradigm of EAE experiments, where ICV infusion into the CNS of either DHF or the DMSO vehicle commenced on Day 13 after the onset of tail weakness. (b) There was no change (p=0.46) in the mean clinical score of mice that received either DHF or DMSO (n=4-5/group, mean ± SD plotted). (c) Longitudinal section of the lateral white matter of the lumbar spinal cord of a sham-EAE mouse at Day 35, showing intact compact myelin as detected by SCoRe imaging (cyan) and fluromyelin staining (white), as compared to spinal cord from a MOG-EAE mouse at Day 35 that received DHF, where there remained prevalent myelin debris. Scale bar = 50μm.

## Discussion

We sought to determine if direct infusion into the CNS of the TrkB agonist DHF, could promote myelin repair in two animal models used preclinically to evaluate therapeutics for MS. We found that DHF failed to promote myelin repair in the cuprizone model of de- and remyelination, with no increase in the proportion of myelinated axons after cuprizone withdrawal compared to the aCSF treated vehicle. This was despite increases in the percentage area of the caudal corpus callosum positive for MBP staining. DHF also failed to alter the clinical course of EAE, an immune-driven model of demyelination, when it was administered via ICV infusion after the initial onset of tail and/or hindlimb weakness. These results are inconsistent with previous findings, wherein recombinant BDNF, and other compounds, TDP6 and LM22A-4, shown to activate TrkB signalling in the cuprizone model, and in multiple *in vitro* contexts [12,14,21,22], did promote remyelination by increasing the proportion of myelinated axons and/or density of post-mitotic oligodendrocytes [13,14]. They also contrast findings in EAE, where intraperitoneal administration of DHF improved the clinical course, preserved myelination and reduced neuroinflammation in the spinal cord [15]. In the current study, the only indication that DHF may have pro-myelinating properties was a significant increase in OPC density in the caudal corpus callosum at the end of 7 days of remyelination.

Proliferation of OPCs is required to replace the myelinating oligodendrocytes that are killed during cuprizone induced demyelination. The increased OPC density in response to DHF infusion during the remyelinating phase in cuprizone-demyelinated mice could be a priming event that precedes robust myelin repair that extends past the 7-day treatment period examined here. ICV infusion of LM22A-4 also significantly increased OPC density in the caudal corpus callosum after 7-days of infusion in cuprizone-demyelinated mice [14]. However, the LM22A-4-mediated increase in OPC density was accompanied by a strong pro-differentiation effect, typical of oligodendroglial TrkB signalling [23], with a significant increase in post-mitotic oligodendrocytes and myelinated axons [14]. Promoting oligodendrocyte differentiation and myelination of axons are two measures of effective myelin repair that DHF was unable to elicit in a mouse model where endogenous remyelination commences within 3-days [13,14,24,25]. More importantly, these are capabilities that are absolutely essential for a candidate remyelinating therapy for MS, in which the primary aim is to overcome arrested OPC differentiation [6,26].

The poor efficacy of DHF as a remyelinating therapeutic for MS when administered to the CNS was also demonstrated in EAE as it did not modify the clinical course, and demyelination and neuroinflammation remained extensive. Testing potential MS therapeutics in EAE is widely considered to be required due to the complex interactions between the immune system and the CNS that occur in MS, and most disease-modifying MS treatments clinically available are immuno-modulatory and have been tested in this model [7]. Previously, DHF was shown to prevent EAE from progressing beyond tail weakness when administered via intraperitoneal injection from the day of EAE induction [15]. This is in contrast with the strategy used here, where DHF was administered directly to the CNS via ICV infusion and did not commence until EAE mice demonstrated tail and/or hindlimb weakness. The timing of treatment from the onset of clinical signs is more likely to represent what MS patients will experience.

The dose of DHF used here is 131x lower than the intraperitoneal delivered dose [15] and this may account for the lack of efficacy in the current study. However, it is reflective of the same scaling used of effective doses of LM22A-4 and TDP6 from *in vitro* [12,27] to *in vivo* application in the cuprizone model (13,14). It is also possible that the ICV infusion of DHF into the cerebrospinal fluid failed to reach the lumbar and sacral spinal cord which are severely affected by peripheral immune cell infiltration and demyelination that precipitate the ascending paralysis characteristic of EAE. This seems highly likely based on the extent of demyelination present in the lumbar spinal cord of DHF-treated EAE mice at Day 35. It would be interesting to compare how high-dose peripheral delivery of DHF from clinical onset influences the clinical course of EAE, and if this strategy would be as effective as the prophylactic regime previously tested [15].

Alternatively, it is possible that DHF mediates its effects in a TrkB-independent manner or with altered timing that is not directly comparable to typical BDNF-TrkB signalling dynamics. Increased OPC density in cuprizone remyelination and the reduced severity of EAE in the previous study [15] in response to DHF treatment are not what would be expected based on the effect of TrkB activation on autoreactive T-cells [28] or oligodendroglia [5,13,14,23]. TrkB activation on autoreactive T-cells has been reported to promote their survival and propagate the autoimmune attack against myelin [28]. Instead, peripheral administration of DHF from the day of MOG sensitisation exerted an anti-inflammatory effect with significantly fewer CD45+ leukocytes, CD3+ T-cells, CD20+ B-cells and CD11b+ macrophages present in the spinal cord of the peripherally DHF-treated EAE mice [15]. It is possible this response could been due to the anti-oxidant properties of DHF [17,18,29] which are able to promote neuronal cell line survival in a TrkB-independent manner[17,18]. Similarly, oligodendroglial TrkB activation is strongly associated with a pro-differentiation effect mediated by MAPK-ERK1/2 signalling [5,30]. Interestingly, DHF has been reported to be biased towards PI3K-Akt signalling [16,31] which is associated with promoting OPC survival [32,33]. This could potentially explain the increase in OPCs. Further, DHF, like LM22A-4 and 6 other TrkB agonists, have come under scrutiny for their inability to transduce TrkB signalling in a BDNF-like manner [34]. Overall, the data from DHF-treated cuprizone mice, the previous study in EAE [15], and several other studies examining the mode of action of DHF [16–18,29,34] suggest that DHF may not act in a manner consistent with typical BDNF-TrkB signalling. This indicates that studies using DHF to mimic of BDNF activation of TrkB to elucidate TrkB receptor function should be interpreted with this caveat in mind.

The lack of pro-myelinating effect of DHF after cuprizone induced demyelination and its failure to modify the disease course of EAE when directly administered to the CNS suggest that DHF does not have the key characteristics required to be a viable remyelinating therapeutic for demyelinating diseases. Further, the results of the current study, combined with existing literature on DHF and its mode of action, suggest that DHF may not always elicit its effects in a BDNF-like manner.

## Materials & Methods

### Experimental animals

Male and female C57BL/6 mice, aged 6-7-weeks old were purchased from the Animal Resource Centre (Perth, Australia) and group housed in specific pathogen-free conditions at the Melbourne Brain Centre Animal Facility, under a 12 hour light / dark cycle with *ad libitum* access to food and water. Animals were habituated to the new housing environment for a minimum of 7 days. All procedures were performed with approval from the Florey Institute of Mental Health and Neuroscience Animal Ethics Committee (#12-042, #18-024) and followed the Australian Code of Practice for the Care and Use of Animals for Scientific Purposes.

### Cuprizone-induced demyelination

At 8 weeks of age, female mice were fed 0.2% cuprizone in normal chow (Teklad Custom Diets) for 6-weeks to induce demyelination. Cuprizone was then removed, and mice were killed (*n*=2/cohort) or underwent intracerebroventricular pump implantation to receive aCSF, DMSO vehicle or DHF for 7 days as previously described. Only female mice were used in cuprizone experiments due to the increased incidence of MS in females [35].

### Experimental autoimmune encephalomyelitis and clinical monitoring

To induce experimental autoimmune encephalomyelitis (EAE), male and female mice, aged 8 weeks received subcutaneous injections to both flanks and base of the tail of MOG_35-55_ peptide (125μg total, Mimotopes) emulsified 1:1 (vol/vol) in Freund’s complete adjuvant (BD Cat# 263810) containing additional *Mycobacterium tuberculosis* (5mg/mL, BD Cat# 231141). A subset of mice received a sham emulsion of 1:1 phosphate buffered saline (PBS) in Freund’s complete adjuvant with 5mg/mL *M. tuberculosis*. Injections of emulsion were followed by intraperitoneal injection of 400ng pertussis toxin (List Biological Laboratories, Sapphire Biosciences Cat# 70323-44-3) in 0.1mL in sterile PBS. Mice were anaesthetized for all injections on Day 0 with exposure to 4% isoflurane in air which was maintained for less than 5 mins at 0.5-1.5% isoflurane through a nose cone. On Day 3, all mice received a second intraperitoneal injection of 400ng pertussis toxin. Mice were weighed and assessed for EAE clinical signs daily. EAE clinical scores was graded according to Table 1.

**Table 1:** Scoring criteria for daily assessment of clinical signs in EAE mice.

| Score | Description |
| --- | --- |
| 0 | Normal appearance, tail lifts and normal gait. |
| 1 | Normal gait, but tail is weak and is slow or fails to curl around finger when stroked. |
| 1.5 | Tail is weak, is slow / fails to curl around finger when stroked. Normal appearing gait, but some hindlimb weakness when performing hindlimb clasp test [39]. |
| 2.0 | Complete tail paralysis, can no longer be lifted by mouse and fails to curl around finger when stroked. |
| 2.25 | Complete tail paralysis and some hindlimb weakness when performing hindlimb clasp test. |
| 2.5 | Complete tail paralysis and hindlimb weakness when performing hindlimb clasp test. |
| 2.75 | Complete tail paralysis and paralysis of one hindlimb when walking. |
| 3.0 | Complete tail and hindlimb paralysis, but maintains righting reflex, when placed on back / side. |
| 3.5 | Complete tail and hindlimb paralysis, with forelimb weakness and loss of righting reflex or signs of urinary / faecal incontinence. Requires immediate euthanasia. |
| 4.0 | Spontaneous death. |

### Intracerebroventricular delivery of 7,8-dihydroxyflavone (DHF)

Cuprizone and EAE mice received intracerebroventricular (ICV) osmotic minipumps (Azlet) vehicle as previously described [13] of 762ng/day 7,8-dihydroxyflavone (equivalent to 250μM for the 7 day infusion with 0.5μL/hr flowrate, or 500μM for the 21 day infusion with 0.25μL/hr flowrate, DHF), artificial aCSF (cuprizone mice only) or 14% (v/v) DMSO. DHF doses were based on previous *in vitro* studies, wherein 250 to 500nM of DHF was required to reach the same level of neuronal survival as BDNF [11]. Briefly, mice were anaesthetised with 2-4% isoflurane in normal air, scalp was shaved, and mice were fitted into a stereotaxic frame receiving 1.5-2.5% isoflurane in air via a nose cone. A sagittal incision was made across the midline of the scalp, and a 1.5mm burr hole was made at +0.5mm rostral and −0.7mm lateral from Bregma using a microdrill. Reservoirs were placed subcutaneously along the flank and cannulae were implanted and fixed in place using an ethyl cyanoacrylate adhesive (Loctite). Vicyrl sutures (Ethicon, Cat# J492G) were used to close head wounds and animals placed in recovery chamber at 32°C and closely monitored until ambulatory, before being single-housed in standard cages (Technoplast). Cuprizone treated mice received 7-day pumps (Azlet #1007D) on the final day of cuprizone exposure before being returned to normal chow and tolerated the procedure well. All ICV minipumps were implanted in cuprizone treated animals on the same days as animals reported in (13, 14). EAE mice received 28-day pumps (Azlet #2004) on Day 13 if they demonstrated tail weakness or tail paralysis. Of 19 EAE mice that underwent the procedure 10 died due to anaesthetic complications or were excluded due to poor wound healing. One EAE-induced mouse did not develop clinical signs by Day 13 and was excluded. Due to high mortality and exclusion rate, ICV implantation in EAE mice was discontinued and available sample size is limited.

### Tissue collection, processing, and immunofluorescence

All mice were anaesthetised with isoflurane and euthanased with barbiturates (Lethabarb, Vibrac Australia) before transcardial perfusion with sterile 0.1M PBS, then ice-cold 4% paraformaldehyde (PFA). Brains from cuprizone mice and the lumbar enlargement of spinal cords from EAE mice were removed. Tissues were post-fixed overnight in 4% PFA, then washed in 0.1M PBS and cryoprotected in 30% sucrose. For electron microscopy (EM) studies of cuprizone mice, the first millimeter of the right hemisphere from the sagittal midline was selected and placed in Kanovsky’s buffer overnight and washed in 0.1M sodium cacodylate before resin embedding at the Centre for Advanced Histology and Microscopy at the Peter MacCallum Centre. Remaining cuprizone brain tissue was oriented sagittally, while EAE spinal cords were oriented longitudinally for embedding in OCT (Tissue-tek) and frozen in isopentane over dry ice. Tissue blocks were stored at −80°C until sectioning at 16μm (brain) or 20μm (spinal cord) using a cryostat maintained at −16 to −17°C, with sections collected on Superfrost^+^ slides. Slides were air-dried and stored at −80°C before use. Approximately 70-100 μm separated adjacent sections on the same slide and sagittal brain sections cut beyond ± 2.64mm from the lateral midline were excluded as previously [13,14].

Primary antibodies for immunostaining are listed in Table 2. Slides were immersed in three exchanges of 0.1M PBS, before overnight incubation at room temperature with primary antibodies diluted in 10% normal donkey serum (NDS) with 0.3% Triton-X 100 in 0.1M PBS. Next, slides were washed three times in 0.1M PBS before incubation with appropriate AlexaFluor-conjugated secondary antibodies (ThermoFisher) for 2 hours at room temperature, after which slides underwent a final wash and were cover slipped using aqueous mounting solution (Dako). After air-drying slides were stored at 4°C until imaging. Modifications for MBP staining included immersing slides in ice-cold 100% methanol for 10mins prior first wash. When used, Hoeschst33342 (ThermoFisher) was the nuclear counterstain.

**Table 2:**
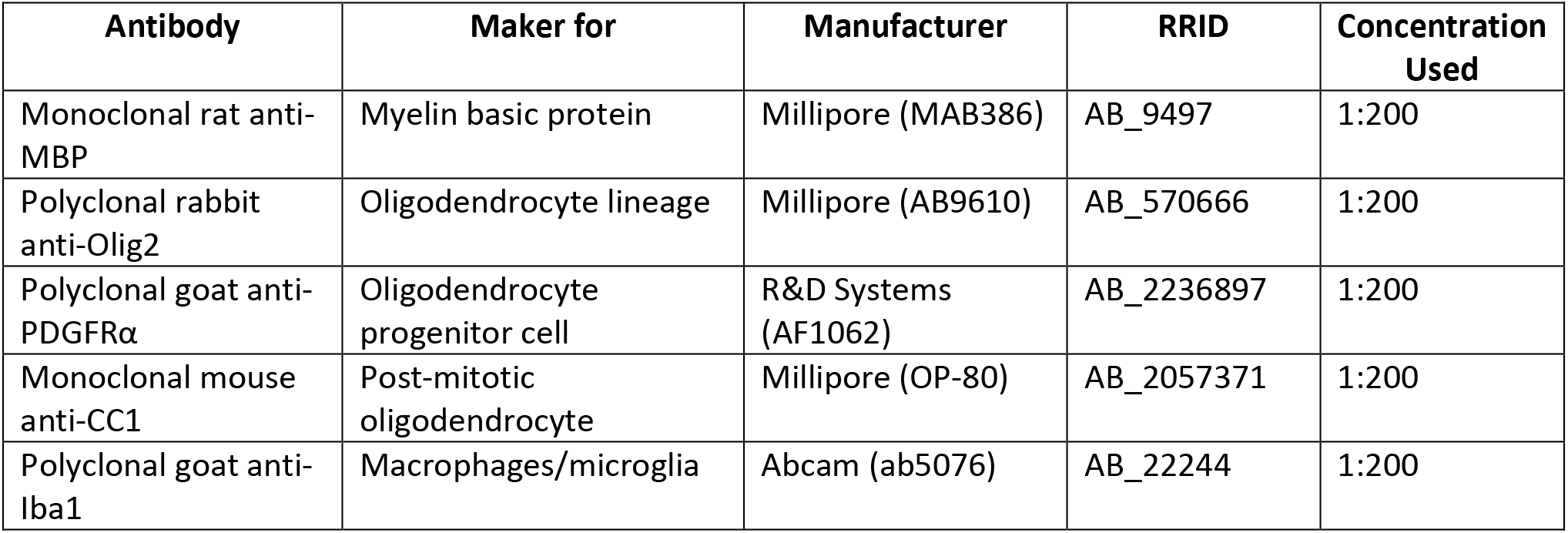
Antibodies used for immunostaining

| Antibody | Maker for | Manufacturer | RRID | Concentration Used |
| --- | --- | --- | --- | --- |
| Monoclonal rat anti-MBP | Myelin basic protein | Millipore (MAB386) | AB_9497 | 1:200 |
| Polyclonal rabbit anti-Olig2 | Oligodendrocyte lineage | Millipore (AB9610) | AB_570666 | 1:200 |
| Polyclonal goat anti-PDGFRα | Oligodendrocyte progenitor cell | R&D Systems (AF1062) | AB_2236897 | 1:200 |
| Monoclonal mouse anti-CC1 | Post-mitotic oligodendrocyte | Millipore (OP-80) | AB_2057371 | 1:200 |
| Polyclonal goat anti-Iba1 | Macrophages/microglia | Abcam (ab5076) | AB_22244 | 1:200 |

### Fluorescence imaging and analysis

All imaging was performed blinded to treatment group. For sagittal sections, imaging was restricted to the caudal region of the corpus callosum approximately −1.1mm to −3.0mm from Bregma, and tracts contributing to the dorsal hippocampal commissure were excluded as previously [13]. Spinal cord sections were from approximately L2 to S1, with imaging restricted to the lateral white matter tracts. For each analysis, a minimum of three sections per animal were imaged.

MBP staining was imaged under a 20x (0.8 NA) lens using an RGB camera (AxioVision Hr, Zeiss) attached to an epifluorescence Axioplan2 Imager (Zeiss) microscope using consistent exposure times. All remaining stains were imaged using Zeiss LSM510 Meta, LSM780 or LSM880 confocal microscope with 405nm, 488nm, 561nm or 633nm laser lines. Images acquired on the LMS780 and 880 were processed (tiles stitched, and maximum z-projections produced) using Zen Black software. Uniform acquisition settings were used for each staining batch. Brightness and contrast were adjusted if needed for counting and presentation purposes.

Oligodendroglial cell counts were performed manually as described in [14] using FIIJ/Image J and expressed as number of cells/mm^2^.

### Electron microscopy and analysis

Electron microscopy and analysis was performed as described in [13]. Briefly, after region selection on semi-thin sections collected on glass slides and stained with 1% toluidine blue, ultrathin sections were collected on 3mm x 3mm copper grids and examined using a JEOL 1011 transmission electron microscope, and imaged using a MegaView III CCD cooled camera operated with iTEM AnalySIS software (Olympus Soft Imaging Systems). A minimum of six distinct fields of view were captured at 10 000x magnification per animal, and used to count myelinated axons in FIJI/Image J. For this analysis, DHF treated animals were compared to animals that solely received aCSF vehicle treated animals previously published in [13,14] as DMSO treated animals were not processed for EM analyses. Sectioning, post-staining and EM imaging were all performed at the Centre for Advanced Histology and Microscopy, Peter MacCallum Centre.

### Spectral confocal reflectance (SCoRe) microscopy

SCoRe microscopy on longitudinally oriented spinal cord from EAE mice was performed as described in [36], and combined with fluoromyelin (ThermoFisher Cat# F34652, RRID: AB_2572213) and Iba1 immunostaining. Briefly, before imaging sections were washed in 0.1M PBS and incubated at 4°C with goat-anti Iba1 (Table 2) in 10% NDS without 100% Triton-X 100 to keep myelin membranes intact for 2 nights in a humified chamber. Sections were washed and incubated with donkey anti-goat AlexaFluor488 for 4 hours at room temperature, before being washed and then incubated in 50% fluoromyelin in PBS for 20 min, and washed thoroughly, before being mounted with #1.5 coverslips (Zeiss, 1.5H, 170±5μm). SCoRe images were captured using a Zeiss LSM880 confocal microscope using 40x oil immersion lens (1.3 NA) and immersion oil (Zeiss, Immersol 518F), with a refractive index of 1.518. Closely, matched refractive index of the coverslip and immersion oil prevented reflection of light from the glass coverslip interfering with SCoRe signal. Lasers with 488nm, 561nm and 633nm wavelengths were passed through acousto-optical tunable filters (AOTF) 488-640 filter/splitter and a 20/80 partially reflective mirror. Reflected light was collected using three photodetectors set to collect light through narrow bands defined by prism and mirror slides, centred around the laser wavelengths.

### Experimental design and statistical analyses

All quantitative assessments were performed by an observer blinded to animal and treatment identity with a minimum of 3 animals per group assessed. All EAE clinical score assessments were performed by observers blinded to treatment group. Statistical testing including 1-way ANOVA and student’s unpaired t-tests were performed in GraphPad Prism (v.8). Restriction maximum likelihood (REML) mixed models were performed in jamovi [37] using GAMLj (v.2.0.6) to evaluate the effect of treatment and day post-induction on clinical score [38]. A p-value less than 0.05 was considered significant. Mean ± standard deviation (SD) plotted in all graphs.

## Acknowledgements

We thank Dr Chris Dwyer and Michele Binder (Florey Institute for Neuroscience and Mental Health) for their assistance with the EAE model. We acknowledge the staff and facilities of the Biological Optical Microscopy Platform (University of Melbourne), Centre for Advanced Histology and Microscopy (Peter MacCallum Centre) and Florey Advanced Microscopy and Immunohistochemistry Service. JLF was supported by a MS Research Australia Postdoctoral Fellowship (#14-056). This work was funded by an Australian National Health and Medical Research Council Project Grant (#APP1105108) awarded to SSM.

This work was supported by the Assistant Secretary of Defense for Health Affairs, endorsed by the Department of Defense, USA, through the Multiple Sclerosis Research Program W81XWH-17-MSRP-IIRA under award no. MS170031 to Simon Murray. Opinions, interpretations, conclusions, and recommendations are those of the authors and not necessarily endorsed by the Assistant Secretary of Defense for Health Affairs or Department of Defense (USA).

## Author contributions

JLF: conceptualization, data curation, formal analysis, investigation, project administration, supervision, visualization, writing – original draft preparation, writing – review & editing. RJW: investigation, methodology, project administration, validation, writing – revieW & editing. ARP, RO & OE: investigation. DGG: methodology, resources, validation, writing – review & editing. SSM: conceptualization, funding acquisition, project administration, resources, supervision, writing – review & editing.

## Data Availability Statement

Data will be made available upon request to the corresponding authors.

## Supplementary Figure

**Fig. S1:**
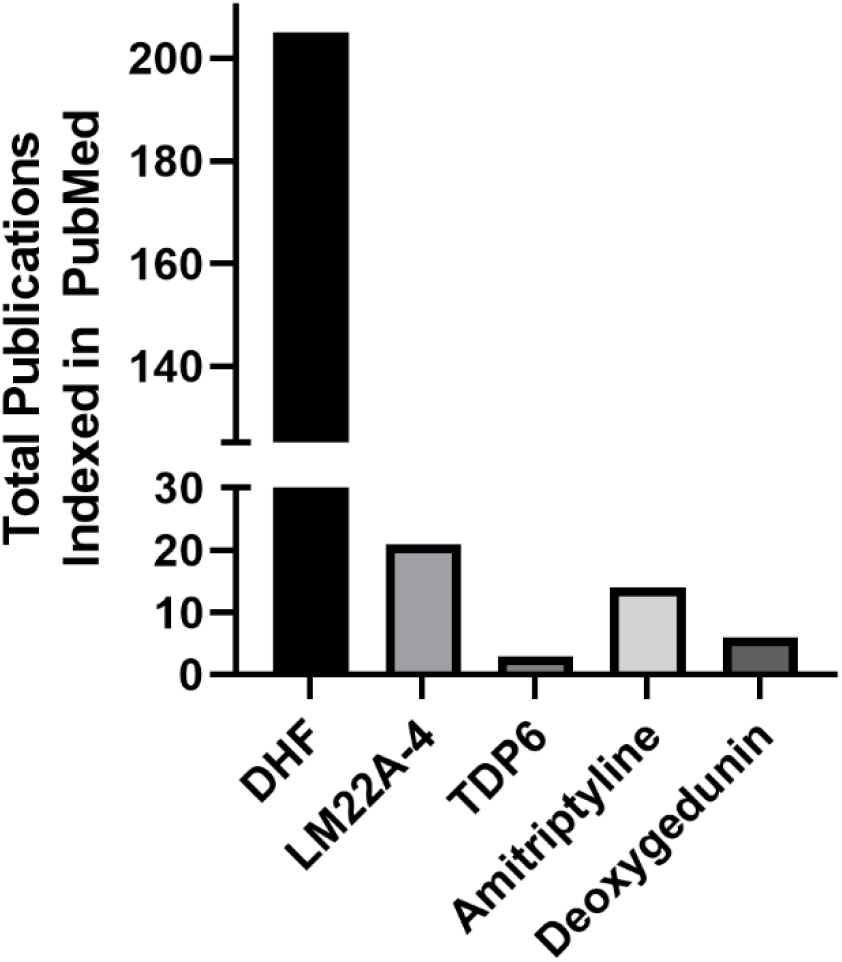
Total number of publications for each TrkB agonist indexed in PubMed. Each TrkB agonist was used as a search term in PubMed and the ‘results by year’ were downloaded and summed. Note for DHF and Amitriptyline TrkB was included as an additional search term and only publications since their characterisation as partial TrkB agonists were used (ie. from 2010 for DHF and 2007 for amitriptyline). Search date: 7 October 2022

